# Cryo-EM structures of Aβ40 filaments from the leptomeninges of individuals with Alzheimer’s disease and cerebral amyloid angiopathy

**DOI:** 10.1101/2023.10.05.561069

**Authors:** Yang Yang, Alexey G. Murzin, Sew Peak-Chew, Catarina Franco, Kathy L. Newell, Bernardino Ghetti, Michel Goedert, Sjors H.W. Scheres

## Abstract

We used electron cryo-microscopy (cryo-EM) to determine the structures of Aβ40 filaments from the leptomeninges of individuals with Alzheimer’s disease and cerebral amyloid angiopathy. In agreement with previously reported structures, which were solved to a resolution of 4.4 Å, we found three types of filaments. However, our new structures, solved to a resolution of 2.4 Å resolution, revealed differences in the sequence assignment that redefine the fold of Aβ40 peptides and their interactions. Filaments are made of pairs of protofilaments, the ordered core of which comprises D1-G38. The different filament types comprise one, two or three protofilament pairs. In each pair, residues H14-G37 of both protofilaments adopt an extended conformation and pack against each other in an anti-parallel fashion, held together by hydrophobic interactions and hydrogen bonds between main chains and side chains. Residues D1-H13 fold back on the adjacent parts of their own chains through both polar and non-polar interactions. There are also several additional densities of unknown identity. Sarkosyl extraction and aqueous extraction gave the same structures. By cryo-EM, parenchymal deposits of Aβ42 and blood vessel deposits of Aβ40 have distinct structures, supporting the view that Alzheimer’s disease and cerebral amyloid angiopathy are different Aβ proteinopathies.

## INTRODUCTION

Alzheimer’s disease (AD) is defined neuropathologically by the presence of abundant filamentous plaques and tangles in the brain parenchyma, with the characteristics of amyloid [1]. Plaques are made predominantly of Aβ42, whereas tangles consist of hyperphosphorylated microtubule-associated protein tau. In addition, many individuals with AD have deposits of Aβ40 in leptomeningeal and parenchymal blood vessels, chiefly small arterioles and capillaries, giving rise to Aβ cerebral amyloid angiopathy (CAA) [2,3]. CAA is the accumulation of amyloidogenic proteins, most often Aβ40, in cerebral blood vessel walls. Nearly half of end-stage AD patients have moderate-to-severe CAA [4]. Even though they share Aβ deposition, AD and CAA can occur independently [5]. Aβ CAA is mostly sporadic, but it can also be inherited [6,7] or acquired [8,9]. The most common clinical manifestations are lobar intracerebral haemorrhage and cognitive impairment.

In 2019, the cryo-EM structures of Aβ40 filaments from the leptomeninges of three individuals with AD and CAA were reported at a resolution of 4.4 Å [10]. For each individual, three types of filaments were observed. In the smallest filament type, two identical protofilaments were in evidence, with the C-terminus being less well resolved than the rest of the molecule. Four cross-β strands were typical of each protofilament and the filament twist was right-handed. Filament types made of four and six protofilaments were also observed. Here we report the structures of Aβ40 filaments from the leptomeninges of three individuals with AD and CAA, with resolutions up to 2.4 Å. Compared to the previously reported structures [10], our higher resolution structures reveal a difference in the sequence assignment that redefines the filament fold of Aβ40 peptides and their interactions. Using either sarkosyl or water-based methods to extract leptomeningeal Aβ40 filaments led to identical folds.

## MATERIALS AND METHODS

### Cases of Alzheimer’s disease and cerebral amyloid angiopathy

Leptomeninges from three neuropathologically confirmed cases of AD and CAA were peeled off from the region overlying the thawed cerebral cortex, before being frozen a second time. Case 1 was a male who died with sporadic late-onset AD aged 89 years; the duration of illness was 12 years and his brain weight was 1,098 g. Case 2 was a male who died with sporadic early-onset AD aged 61 years; the duration of illness was 6 years and his brain weight was 1,048 g. Case 3 was a female who died with sporadic early-onset AD aged 70 years; the duration of illness was 9 years and her brain weight was 1,380 g.

### Filament extraction

Sarkosyl-insoluble material was extracted from the leptomeninges of cases 1-3, as described [11]. Tissues were homogenised in 20 vol buffer A (10 mM Tris-HCl, pH 7.5, 0.8 M NaCl, 10% sucrose and 1 mM EGTA), brought to 2% sarkosyl and incubated for 30 min at 37° C. The samples were centrifuged at 10,000 g for 10 min, followed by spinning of the supernatants at 100,000 g for 60 min. The pellets were resuspended in 50 μl/g 20 mM Tris-HCl, pH 7.4, 50 mM NaCl and used for cryo-EM. For aqueous extraction, as described previously [10,12], leptomeningeal tissues (0.2-0.5 g) were cut with a scalpel into cubes of less than 1 mm^3^ and washed 4 times with 500 μl Tris-Calcium buffer (20 mM Tris, pH 7.4, 138 mM NaCl, 2 mM CaCl_2_, 0.1% (w/v) NaN_3_, pH 8.0) at 4° C. Following each wash, the samples were centrifuged at 3,000 g for 1 min, with a 5 min spin at 12,000 g after the final wash. The pellets were then incubated overnight with 5 mg/ml *Clostridium histolyticum* collagenase (Sigma-Aldrich) in 1 ml Tris-Calcium buffer at 37° C. Following a 5 min centrifugation at 12,000 g at 4°C, they were washed 5 times with 500 μl 50 mM Tris-HCl, pH 7.4, 10 mM EDTA, with a 5 min centrifugation at 12,000 g after each wash. To collect Aβ filaments, 250 μl cold water was added to the pellets, followed by a 5 min centrifugation at 12,000 g. This step was repeated nine times. The supernatants were combined and used for subsequent experiments.

### Mass spectrometry

Sarkosyl-insoluble pellets were resuspended in 200 μl hexafluoroisopropanol. Following a 3 min sonication at 50% amplitude (QSonica), they were incubated at 37° C for 2 h and centrifuged at 100,000 g for 15 min, before being dried by vacuum centrifugation. Matrix-assisted laser desorption/ionization time of flight (MALDI-TOF) mass spectrometry was carried out.

### Electron cryo-microscopy

Three μl of the sarkosyl-insoluble fractions were applied to glow-discharged (Edwards S150B) holey carbon grids (Quantifoil AuR1.2/1.3, 300 mesh) that were plunge-frozen in liquid ethane using a Vitrobot Mark IV (Thermo Fisher Scientific) at 100% humidity and 4° C. Cryo-EM images were acquired using EPU software on Titan Krios G2 and G4 microscopes (Thermo Fisher Scientific) operated at 300 kV. Images for case 1 were acquired using a Falcon-4i detector (Thermo Fisher Scientific) in electron-event representation mode with a flux of 11 electrons/pixel/s and a Selectris-X energy filter (Thermo Fisher Scientific) with a slit width of 10 eV to remove inelastically scattered electrons. Images for case 2 were acquired using a Falcon-4i detector without energy filter. Images for case 3 were acquired using a Gatan K3 detector in super-resolution counting mode, using a Bio-quantum energy filter (Gatan) with a slit width of 20 eV. Images were recorded with a total dose of 40 electrons per A^2^. See Supplementary Table for further details.

### Helical reconstruction

Datasets were processed in RELION using standard helical reconstruction [13]. Movie frames were gain-corrected, aligned and dose-weighted using RELION’s own motion correction programme [14].

Contrast transfer function (CTF) was estimated using CTFFIND4-1 [15]. Filaments were picked manually. Following 2D classification, polymorphs were identified using the clustering approach FilamentTools (https://github.com/dbli2000/FilamentTools) [16]. Initial models were generated *de novo* from 2D class average images using relion_helix_ inimodel2d [17]. Three-dimensional auto-refinements were performed with optimisation of the helical twist and rise parameters once the resolutions extended beyond 4.7 Å. To improve the resolution, Bayesian polishing and CTF refinement were used to improve resolution [18]. Final maps were sharpened using standard post-processing procedures in RELION and resolution estimates calculated based on the Fourier shell correlation (FSC) between two independently refined half-maps at 0.143 [19] (Supplementary Figure 1).

### Model building and refinement

Atomic models were built manually in the reconstructed cryo-EM densities using Coot [20]. Model refinements were performed using *Servalcat* [21] and REFMAC5 [22,23]. Models were validated with MolProbity [24]. Figures were prepared with ChimeraX [25] and PyMOL [26].

## RESULTS

### Structures of Aβ40 filaments from leptomeninges extracted using sarkosyl

We determined the cryo-EM structures of Aβ filaments from the leptomeninges of three cases of AD and CAA. There were two main types of filaments, the minority type (type 1) comprising a pair of identical protofilaments and the majority type (type 2) comprising two protofilament pairs running side by side. All filaments had a right-handed twist, as established from the densities for backbone oxygen atoms in the cryo-EM maps. Structures from case 1 reached the highest resolution. The structure of filaments made of two pairs of protofilaments were determined at 2.4 Å resolution; the structure of filaments made of one pair of protofilaments was determined at 2.7 Å (Figure 1a; Figure S1).

**Figure 1.**
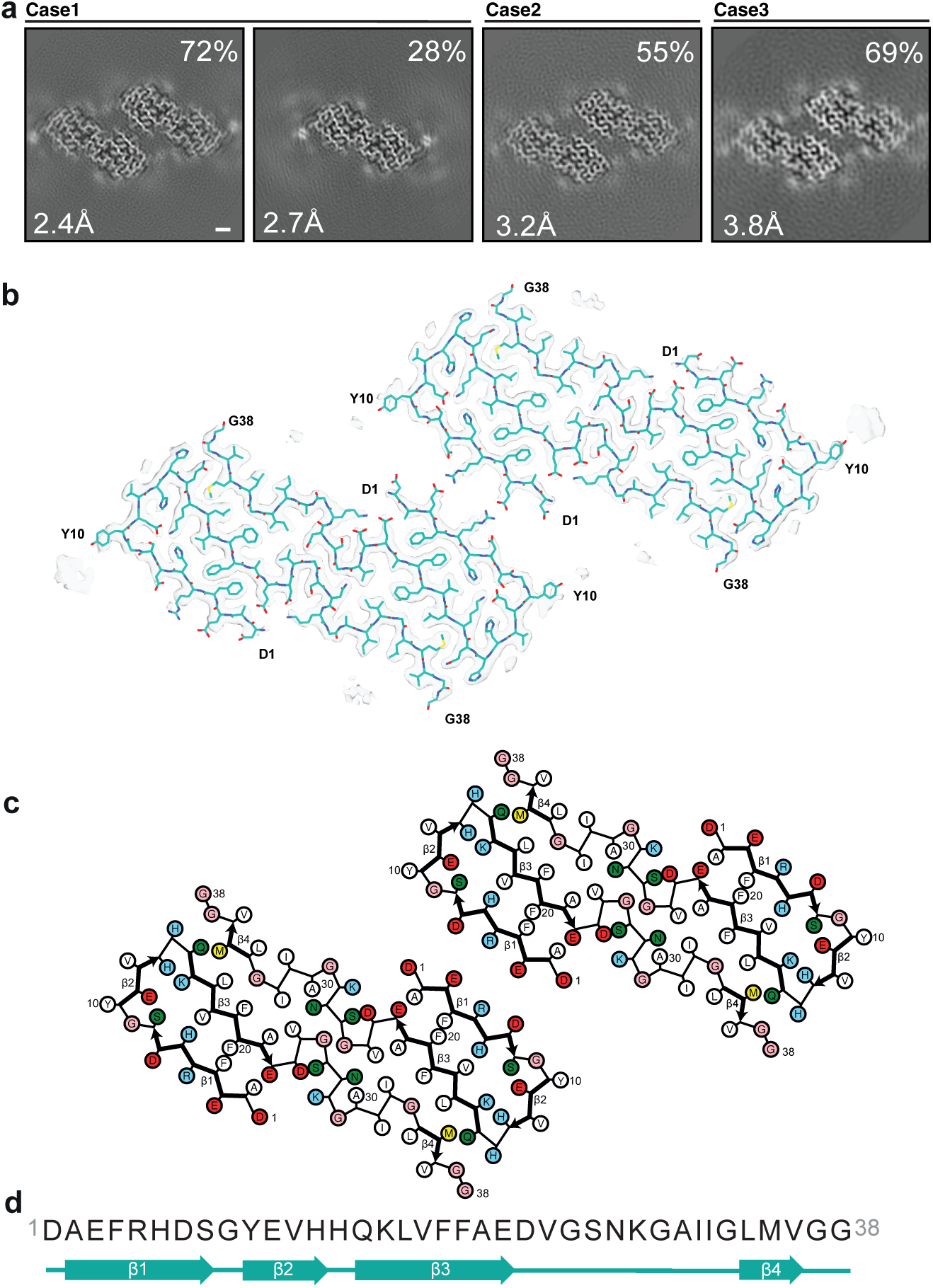
Structures of Aβ40 filaments from leptomeninges extracted using sarkosyl. **a**, Cross-sections of Aβ40 filaments from cases 1-3 perpendicular to the helical axis, with a projected thickness of approximately one rung. Percentages of filaments are shown on the top right. The resolutions of the cryo-EM maps are indicated on the bottom left. Filaments were made of one (type 1) or two (type 2) pairs of protofilaments. Scale bar, 1 nm. **b**, Cryo-EM density maps (transparent grey) and atomic model (cyan) of type 2 Aβ40 filaments with two pairs of protofilaments. **c**, Schematic of the structure shown in panel b. Negatively charged residues are shown in red, positively charged residues in blue, polar residues in green, non-polar residues in white, sulphur-containing residues in yellow and glycines in pink. Thick connecting lines with arrowheads indicate β-strands. **d,** Amino acid sequence of the core of Aβ40 protofilaments, which extends from D1 to G38. Beta-strands (β1-β4) are indicated as thick arrows.

The ordered core of each protofilament consists of residues D1-G38 and comprises four β-strands (β1-β4) that extend from residues 2-8, 10-13, 15-22 and 34-36, respectively (Figure 1b-d). In type 1 filaments, the individual protofilaments pack against each other with 2-start (pseudo-2_1_) helical symmetry through a large interface, made of residues H14-G37 in an extended conformation. This interface is stabilised by hydrophobic interactions of L17, F19, A21, V24, I32 and M35 from both protofilaments and hydrogen bonds from the polar side chains of H14, Q15 and N27 to the main chain of the opposite molecules. The N-terminal segments β1 and β2 fold back on the opposite side of β3, enclosing a network of electrostatic interactions and hydrogen bonds formed by the polar side chains of H6, S8, E11, H13 and K16, and capped by the interlocking aromatic side chains of F4 and F20. The N-terminal substructure formed by the β1-β3 region is similar to that derived from the low-resolution map [10].

Type 2 filaments comprise two pairs of protofilaments that pack against each other with C2 symmetry (Figure 1b). Pairs of leptomeningeal Aβ40 protofilaments interact through salt bridges between E3 from one dimer and R5 from the other dimer, and vice versa. A similar association of dimeric filaments into tetramers via electrostatic interactions of the surface residues K16 and E22 was observed in Aβ42 Type 1b filaments [11].

Besides the density for the ordered core of Aβ filaments, there were additional densities in the cryo-EM reconstruction that could not be explained by Aβ. There was a strong density in front of Y10, which was situated at a right angle turn between β1 and β2, which may correspond to a post-translational modification of this residue and/or an external cofactor. There was another strong density opposite a hydrophobic patch made of I31, L34 and V36 in the C-terminal region. In the previously reoported low-resolution maps [10], there was a disconnected density in the equivalent location that was assigned to the C-terminal residues V39 and V40. The unambiguous sequence assignment of our higher-resolution structures renders such an assignment impossible. In principle, the C-terminal residues of Aβ42 or Aβ43 peptides could reach this disconnected density, but these peptides were uncommon in the cases studied here.

By mass spectrometry, Aβ1-40 showed the highest peak in all three cases (Figure 2), indicating that most filaments were made of Aβ1-40. In case 3, a comparable peak was observed for Aβ1-38, consistent with the ordered filament core extending from D1 to G38.

**Figure 2.**
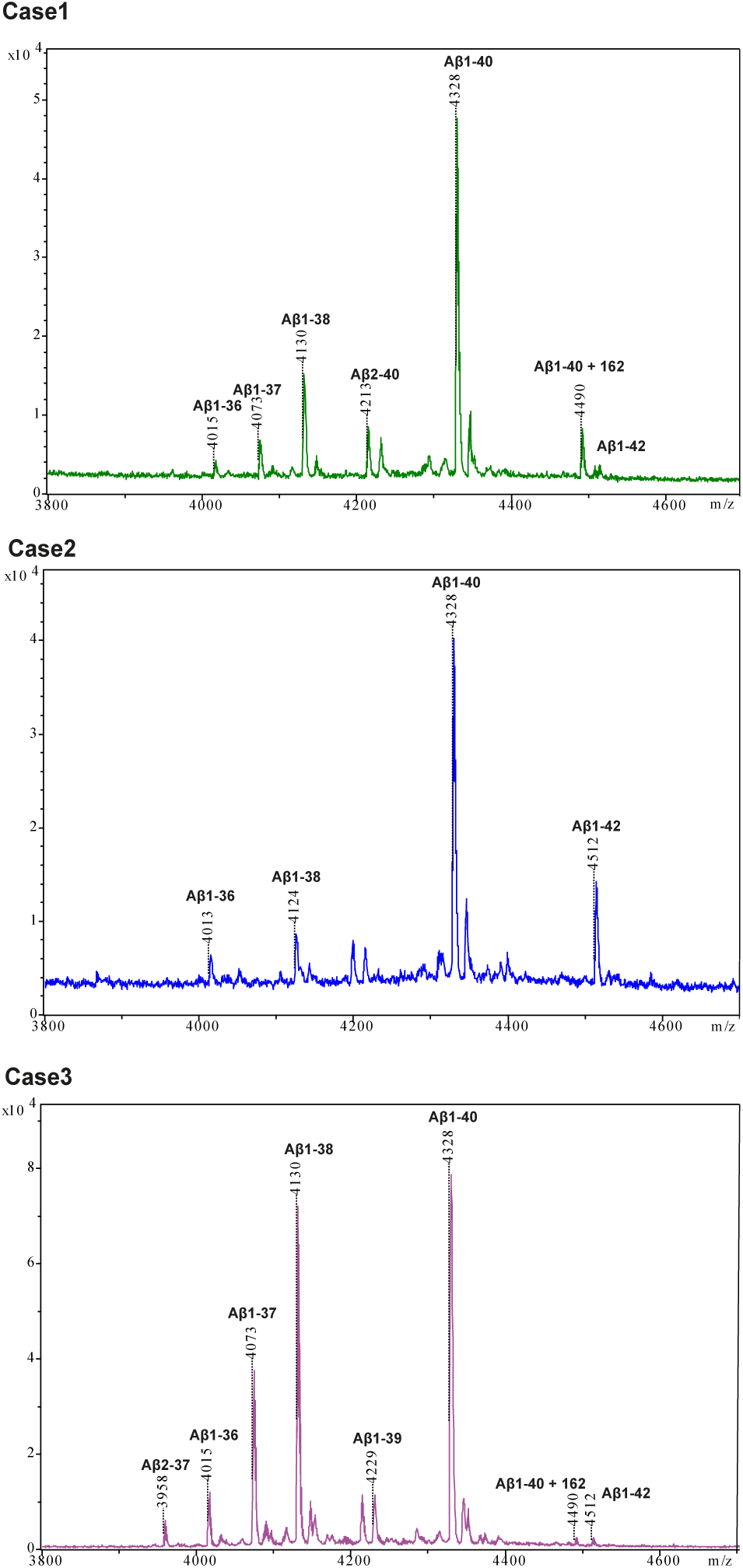
Mass spectrometry. MALDI mass spectra of Aβ from the sarkosyl-insoluble fractions of the leptomeninges of cases 1-3. The peptides with their molecular mass are labelled on top of the spectra.

### Structures of Aβ40 filaments from leptomeninges extracted using aqueous extraction

We also used aqueous extraction of Aβ filaments from the leptomeninges of case 1. Cryo-EM structures of type 1 and type 2 filaments, at 3.1 Å and 2.7 Å resolution, respectively, revealed the same Aβ40 fold as observed using sarkosyl extraction (Figure 3a-c). Aqueous extraction also yielded a third type of filaments with three such pairs of protofilaments (type 3), which was solved at 4.0 Å resolution (Figure 3a). The all-atom root mean square deviation (r.m.s.d.) between protofilaments from type 1 filaments of sarkosyl and aqueous extractions was 1.1 Å. Minor differences were due to a distinct side chain orientation of Y10. The additional density next to this residue was less prominent and more diffuse than in the maps of sarkosyl-extracted filaments, suggesting that a putative cofactor associated with this residue was partially lost during aqueous extraction.

**Figure 3.**
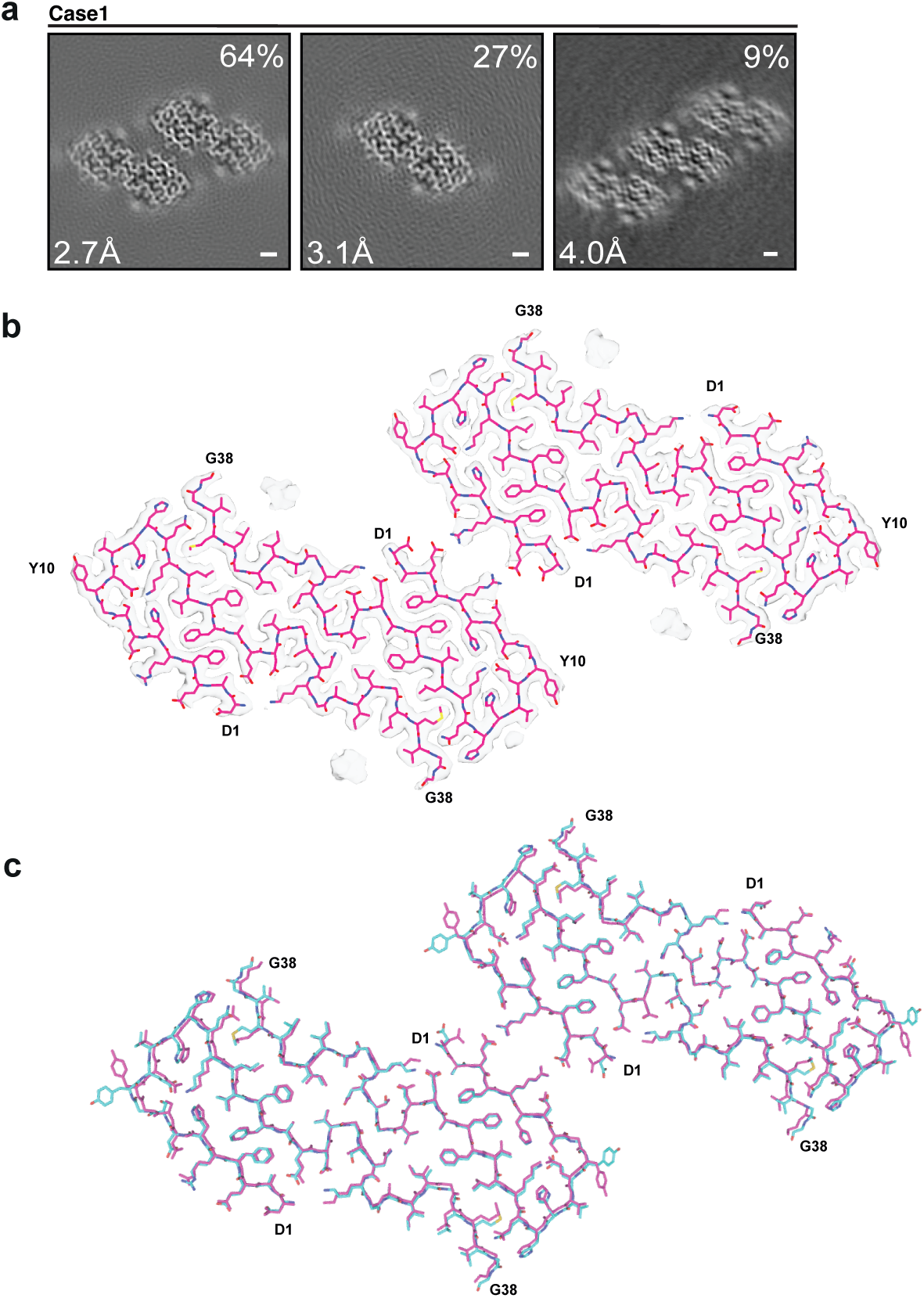
Structures of Aβ40 filaments from leptomeninges extracted using a water-based method. **a**, Cross-sections of Aβ40 filaments from case 1 perpendicular to the helical axis, with a projected thickness of approximately one rung. Percentages of filaments are shown on the top right. The resolutions of the cryo-EM maps are indicated on the bottom left. Filaments were made of one, two or three pairs of protofilaments. Scale bar, 1 nm. **b,** Cryo-EM density maps (transparent grey) and atomic model (pink) of type 2 Aβ40 filaments with two pairs of protofilaments. **c,** Superposition of the structures of type 2 Aβ40 filaments that were extracted using either sarkosyl (cyan) or an aqueous solution (pink).

### Comparison with previous structures

A comparison of our type 1 filament structure at 2.7 Å with the previously reported structure at 4.4 Å [10] revealed differences in the atomic models (Figure 4a,b). The same differences were also present when comparing the previously reported structure of type 1 filaments with the pairs of protofilaments in our 2.4 Å structure of type 2 filaments (not shown). Compared to our model, the previously reported model has the same sequence assignment for the N-terminal 23 residues of Aβ40, but the rest of the sequence is misaligned by up to two residues (Figure 4b,c). Whereas the main-chain assignment is unambiguous in our high-resolution maps, the density in the 4.4 Å map was less clear. It is possible that the misalignment arose from an erroneous assignment of the C-terminal residues of Aβ40 into the disconnected density near the end of the contiguous segment. Fitting our model into the 4.4 Å map (EMD-10204) [10] and comparing it with the deposited model (PDB:6SHS), shows that our model is as good a description of the 4.4 Å map as the deposited model (Figure 4e).

**Figure 4.**
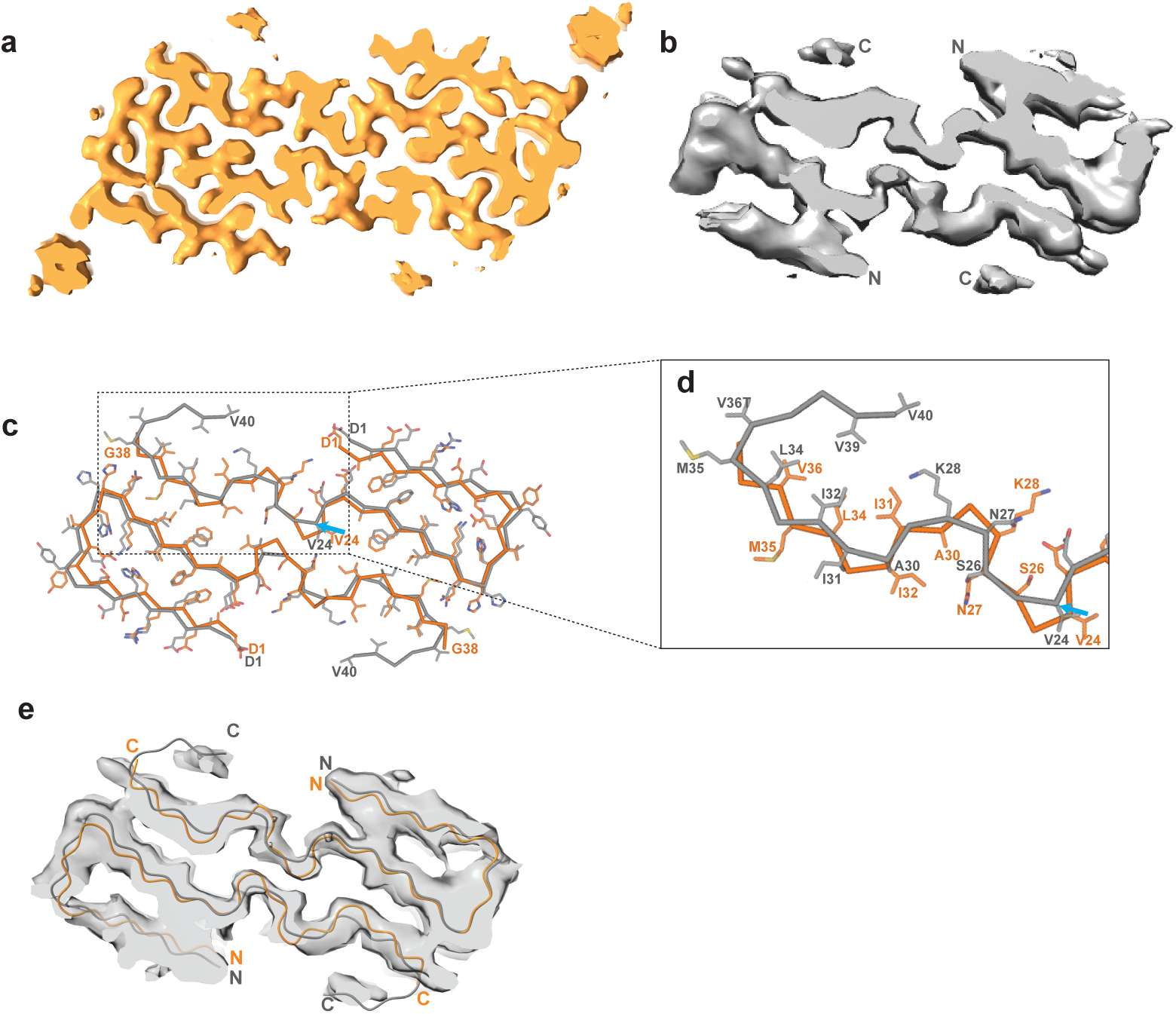
Comparison of the maps and models of Aβ40 filaments described here with those of Kollmer et al [10]. **a,** Map of Aβ40 filaments from leptomeninges with two protofilaments extracted using sarkosyl (present work; in orange). N- and C-termini are indicated. **b,** Map of Aβ40 filaments from leptomeninges with two protofilaments extracted using a water-based method (previous work; in grey). N- and C-termini are indicated. **c,** Atomic models of Aβ40 filaments for the maps shown in panels a and b (present work in orange; previous work in grey). The blue arrow points to V24, from which the two models differ in the register of their main chains. **d,** Zoomed-in view of the structures in panel c. **e,** Atomic model from present work (orange) and from previous work (grey) fitted into the 4.4 Å map from the previous work [10].

## DISCUSSION

We report the cryo-EM structures of Aβ filaments from the leptomeninges of three cases of sporadic AD and CAA with resolutions up to 2.4 Å. In agreement with a previous report [10], we observed three types of right-handed Aβ40 filaments that consisted of one, two or three pairs of protofilaments. Each protofilament extended from D1-G38 and consisted of four β-strands. It follows that Aβ monomers in these filaments have an intact N-terminus and are truncated or disordered for the two C-terminal residues. This contrasts with the structures of Aβ42 filaments from the brains of individuals with AD [11], in which the N-terminal eight or ten residues are disordered and partially truncated, and the C-terminus is ordered. Aβ40 filaments also differ from Aβ42 filaments by the presence of a strong density associated with Y10.

Sarkosyl extraction and aqueous extraction gave nearly identical filament structures, demonstrating that sarkosyl did not influence the structure of Aβ40 filaments. The same was true of Aβ42 filaments and paired helical tau filaments [27]. When using sarkosyl extraction of AD brain tissues, we observed Aβ42 filaments [11]. Only Aβ40 filaments were found when leptomeninges from cases of AD and CAA were extracted with sarkosyl. Both Aβ40 and Aβ42 filaments were found in the brain of a patient with mutation E22G in Aβ (Arctic mutation) following sarkosyl extraction [28].

Our atomic models differ from those reported previously for Aβ40 filaments from the leptomeninges of three individuals with sporadic AD and CAA [10]. Our unambiguous sequence assignment differs from the assignment proposed previously for a low-resolution map [10], thereby redefining the fold of Aβ40 peptides and their interactions. It has been recommended that *de novo* atomic modelling of amyloids should not be performed in RELION reconstructions, when the resolutions are worse than 4.0 Å [17].

Seeded aggregation of amyloid filaments is often used, with the implicit assumption that these filaments have the same structures as the initial seeds. However, this is not necessarily the case, as was shown for seeded aggregation of recombinant α-synuclein with seeds from the putamen of individuals with multiple system atrophy [29]. The cryo-EM structures of synthetic Aβ40 filaments seeded from vascular deposits of brain parenchyma of individuals with CAA have recently been reported [30]. One of the two structures described in this preprint appears to be identical to the structure of the type 1 Aβ40 filaments that we report, including the new sequence assignment of the 15 C-terminal residues.

Without the results described here, this difference in sequence assignment of the seeded structure could have been attributed to a difference between Aβ40 filaments from CAA and CAA+AD, a difference between Aβ40 filaments from leptomeningeal and parenchymal blood vessels, or an artefact of seeded aggregation. However, the present results suggest that the seeds used in [30] contained the same type 1 Aβ40 filaments that we extracted and that their structures were replicated correctly during seeded aggregation.

The cryo-EM structures of Aβ42 filaments from plaques and Aβ40 filaments from blood vessels are different, consistent with the view that AD and CAA are the result of distinct pathogenic processes. For assembled tau and α-synuclein, it has also been shown that specific amyloid structures define different diseases [31]. Medin has been reported to co-aggregate with Aβ40 in blood vessels [32], but we did not observe mixed filaments.

The present findings may have consequences for the development of disease-modifying treatments for AD. Current immunotherapies suffer from common and sometimes serious side effects, also known as amyloid-related imaging abnormalities (ARIAs) [33,34], which appear to be linked to the abundance of pre-existing CAA [35]. Existing antibodies seem to bind to both parenchymal and blood vessel deposits of Aβ. Based on the structural differences, it may be possible to produce conformational Aβ antibodies that bind specifically to either parenchymal or blood vessel deposits. It may also become possible to develop imaging agents that distinguish between parenchymal and blood vessel deposits of Aβ. Accurate knowledge of the atomic structures of the different amyloids will facilitate these developments.

## Acknowledgements

We thank the patients’ families for donating brain tissues. This work was supported by the Electron Microscopy Facility of the MRC Laboratory of Molecular Biology. We thank Jake Grimmett, Toby Darling and Ivan Clayson for help with high-performance computing, as well as Brad Glazier, Urs Kuederli, Rose Richardson and Max Jacobsen for help with neuropathology. We acknowledge Diamond Light Source for access and support of the cryo-EM facilities at the UK’s Electron Bio-Imaging Centre (under proposal bi23268), funded by the Wellcome Trust, the MRC and the Biotechnology and Biological Sciences Research Council (BBSRC). For the purpose of open access, the MRC Laboratory of Molecular Biology has applied a CC BY public copyright licence to any Author Accepted Manuscript version arising.

## Author contributions

KLN and BG identified the patients and performed neuropathology; YY prepared filaments; SPW and CF performed mass spectrometry; YY performed cryo-EM data acquisition; YY, AGM and SHWS performed cryo-EM structure determination; MG and SHWS supervised the project and all authors contributed to the writing of the manuscript.

## Availability of data and materials

Cryo-EM maps have been deposited in the Electron Microscopy Data Bank (EMDB) with the accession numbers EMD-18508 and EMD-18509. Corresponding refined atomic models have been deposited in the Protein Data Bank (PDB) under accession numbers PDB:8QN6 and PDB:8QN7. Please address requests for materials to the corresponding authors.

## SUPPLEMENTARY TABLE

**Table S1.**
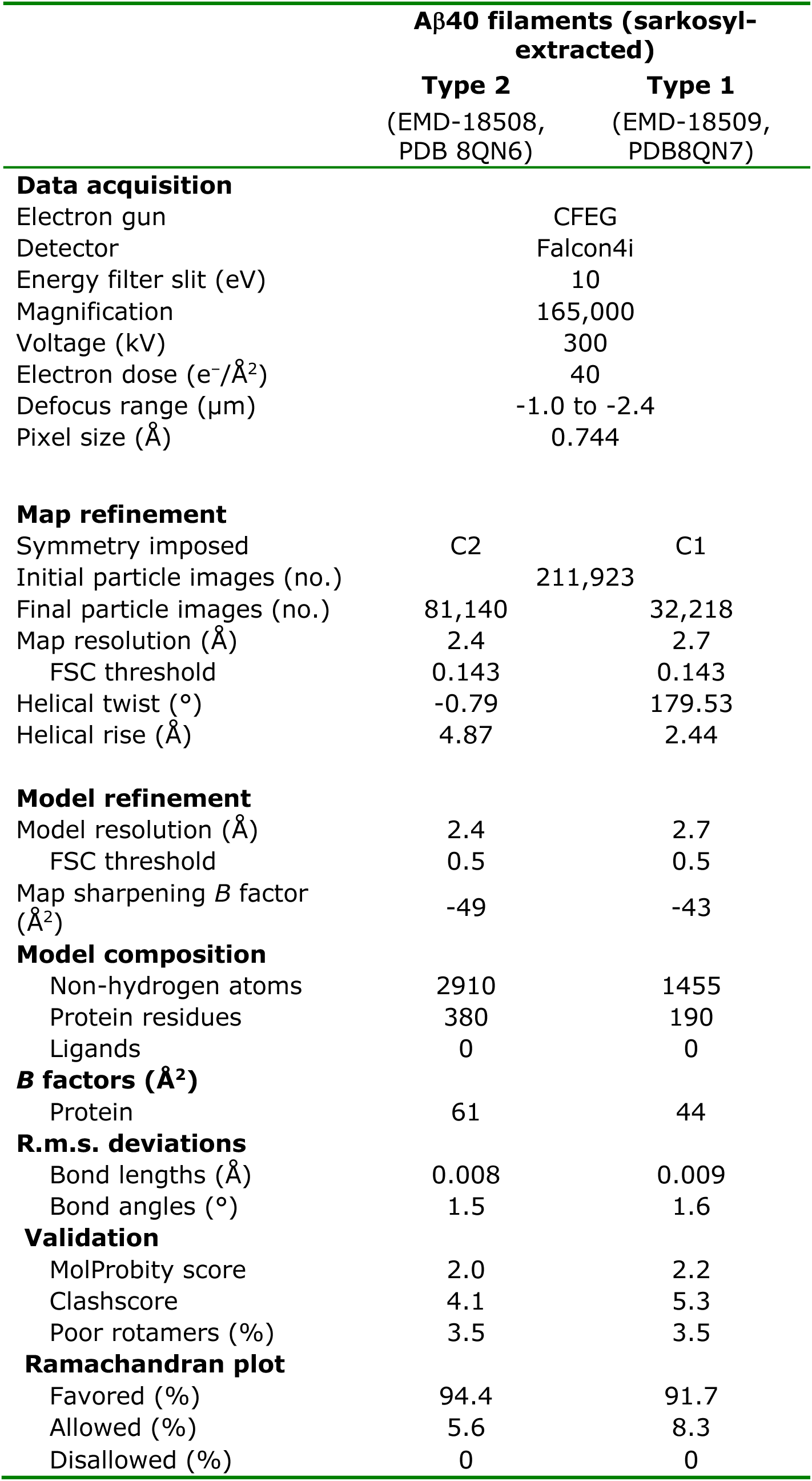
Cryo-EM data acquisition and structure determination.

|  | Aβ40 filaments (sarkosyl-extracted) |  |
| --- | --- | --- |
|  | Type 2<br>(EMD-18508,<br>PDB 8QN6) | Type 1<br>(EMD-18509,<br>PDB8QN7) |
| Data acquisition |  |  |
| Electron gun | CFEG |  |
| Detector | Falcon4i |  |
| Energy filter slit (eV) | 10 |  |
| Magnification | 165,000 |  |
| Voltage (kV) | 300 |  |
| Electron dose (e<sup>-</sup>/Å<sup>2</sup>) | 40 |  |
| Defocus range (μm) | -1.0 to -2.4 |  |
| Pixel size (Å) | 0.744 |  |
| Map refinement |  |  |
| Symmetry imposed | C2 | C1 |
| Initial particle images (no.) | 211,923 |  |
| Final particle images (no.) | 81,140 | 32,218 |
| Map resolution (Å) | 2.4 | 2.7 |
| FSC threshold | 0.143 | 0.143 |
| Helical twist (°) | -0.79 | 179.53 |
| Helical rise (Å) | 4.87 | 2.44 |
| Model refinement |  |  |
| Model resolution (Å) | 2.4 | 2.7 |
| FSC threshold | 0.5 | 0.5 |
| Map sharpening *B* factor (Å<sup>2</sup>) | -49 | -43 |
| Model composition |  |  |
| Non-hydrogen atoms | 2910 | 1455 |
| Protein residues | 380 | 190 |
| Ligands | 0 | 0 |
| *B* factors (Å<sup>2</sup>) |  |  |
| Protein | 61 | 44 |
| R.m.s. deviations |  |  |
| Bond lengths (Å) | 0.008 | 0.009 |
| Bond angles (°) | 1.5 | 1.6 |
| Validation |  |  |
| MolProbity score | 2.0 | 2.2 |
| Clashscore | 4.1 | 5.3 |
| Poor rotamers (%) | 3.5 | 3.5 |
| Ramachandran plot |  |  |
| Favored (%) | 94.4 | 91.7 |
| Allowed (%) | 5.6 | 8.3 |
| Disallowed (%) | 0 | 0 |

## SUPPLEMENTARY FIGURE LEGENDS

**Supplementary Figure 1.**
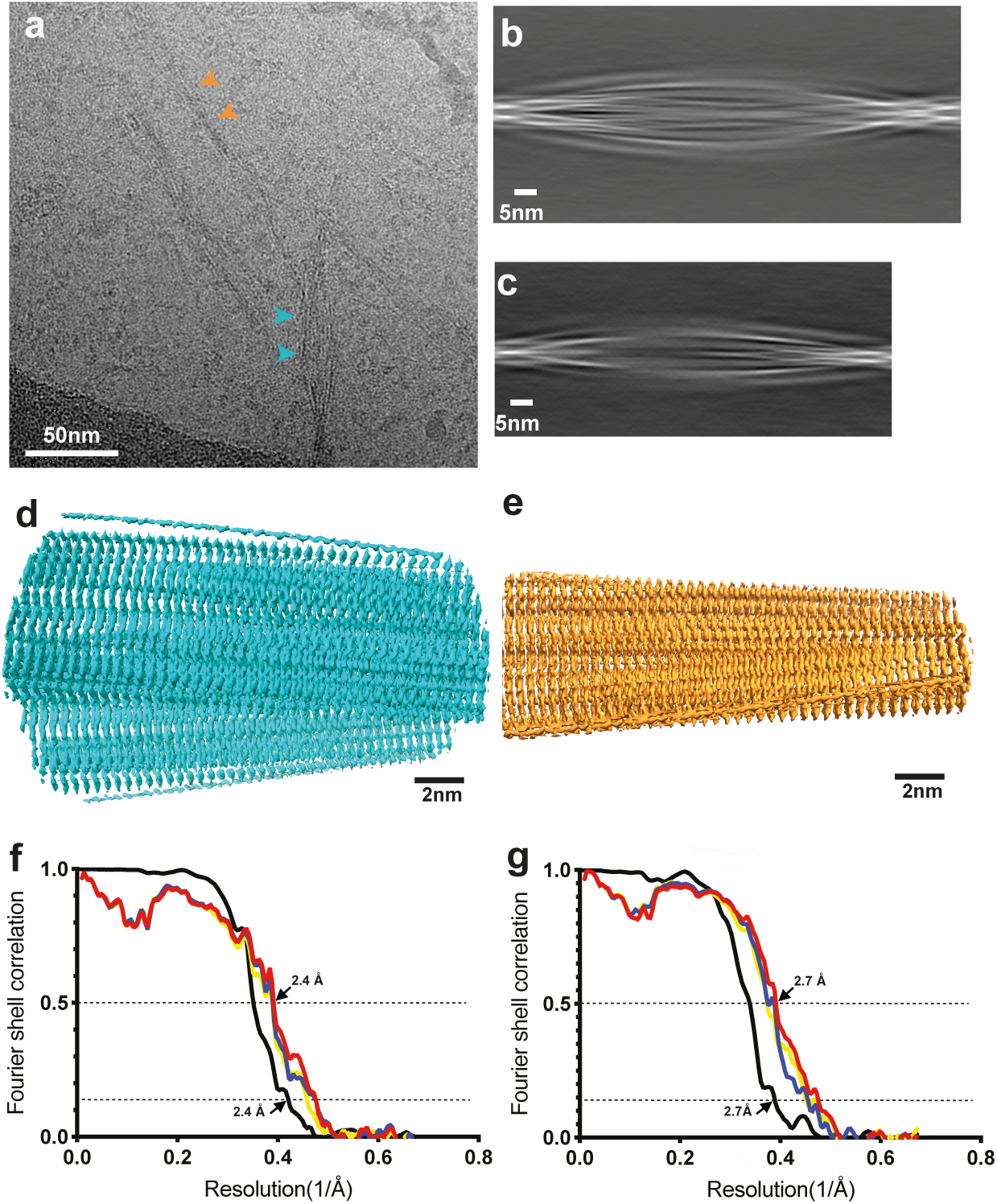
Cryo-EM micrographs and processing details. **a**, Cryo-EM micrographs of Aβ40 filaments from the leptomeninges of case 1 following sarkosyl extraction. Scale bar, 50 nm. **b,c,** 2D class average plots of type 2 (b) and type 1 (c) Aβ40 filaments from the leptomeninges of case 1 following sarkosyl extraction. Scale bar, 5 nm. **d,e,** 3D reconstructions of type 2 (d) and type 1 (e) Aβ40 filaments from the leptomeninges of case 1 following sarkosyl extraction. Scale bar, 2 nm. **f,g,** Fourier shell correlation (FSC) curves for the cryo-EM maps are shown in black; for the refined atomic model against the cryo-EM map in red; for the atomic model refined in the first half map against that half map in blue; for the refined atomic model in the first half map against the other half map in yellow.

## REFERENCES

1. Self WK, Holtzman DM (2023) Emerging diagnostics and therapeutics for Alzheimer disease. Nature Med 29: 2187–2199.

2. Suzuki N, Iwatsubo T, Odaka A, Ishibashi Y, Kitada C, Ihara Y (1994) High tissue content of soluble β1-40 is linked to cerebral amyloid angiopathy. Am J Pathol 145: 452–460.

3. Attems J, Jellinger K, Thal DR, Van Nostrand W (2011) Sporadic cerebral amyloid angiopathy. Neuropathol Appl Neurobiol 37: 75–93.

4. Jäkel L, De Kort AM, Klijn CJM, Schreuder FHBM, Verbeek MM (2022) Prevalence of cerebral amyloid angiopathy: A systematic review and meta-analysis. Alzheimer’s Dement 18: 10–28.

5. Greenberg SM, Bacskai BJ, Hernandez-Guillamon M, Pruzin J, Sperling R, van Veluw SJ (2020) Cerebral amyloid angiopathy and Alzheimer disease – one peptide, two pathways. Nature Rev Neurol 16: 30–42.

6. Levy E, Carman MD, Fernandez-Madrid IJ, Power MD, Lieberburg I, van Duinen SG, Bots GTAM, Luyendijk W, Frangione B (1990) Mutation of the Alzheimer’s disease amyloid gene in hereditary cerebral hemorrhage, Dutch type. Sciencer 248: 1124–1126.

7. Bugiani O, Giaccone G, Rossi G, Mangieri M, Capobianco R, Morbin M, Mazzoleni R, Cupidi C, Marcon G, Giovagnoli A et al (2010) Hereditary cerebral hemorrhage with amyloidosis associated with the E693K mutation of APP. Arch Neurol 67: 987–995.

8. Jaunmuktane Z, Mead S, Ellis M, Wadsworth JD, Nicoll AJ, Kenny J, Launchbury F, Linehan J, Richard-Loendt A, Walker AS et al (2015) Evidence for human transmission of amyloid-β pathology and cerebral amyloid angiopathy. Nature 525: 247–250.

9. Jaunmuktane Z, Quaegebeur A, Taipa R, Viana-Baptista M, Barbosa R, Koriath C, Sciot R, Mead S, Brandner R (2018) Evidence of amyloid-β cerebral amyloid angiopathy transmission through neurosurgery. Acta Neuropathol 135: 671–679.

10. Kollmer M, Close W, Funk L, Rasmussen J, Bsoul A, Schierhorn A, Schmidt M, Sigurdson CJ, Jucker M, Fändrich M (2019) Cryo-EM structure and polymorphism of Aβ amyloid fibrils purified from Alzheimer’s brain tissue. Nature Commun 10: 4760.

11. Yang Y, Arseni D, Zhang W, Huang M, Lövestam S, Schweighauser M, Kotecha A, Murzin AG, Peak-Chew SY, Macdonald J et al (2022) Cryo-EM structures of amyloid-β 42 filaments from human brains. Science 14: 167–172.

12. Annamalai K, Gührs KH, Koehler R, Schmidt M, Michel H, Loos C, Gaffney PM, Sigurdson CJ, Hegenbart U, Schönland S et al (2016) Polymorphism of amyloid fibrils in vivo. Angew Chem Int Ed 55: 4822–4825.

13. He S, Scheres SHW (2017) Helical reconstruction in RELION. J Strucct Biol 198: 163–176.

14. Zivanov J, Nakane T, Forsberg BO, Kimanius D, Wagen WJ, Lindahl E, Scheres SHW (2018) New tools for automated high-resolution cryo-EM structure determination in RELION-3. eLiife 7: e42166.

15. Rohou A, Grigorieff N (2015) CTFFIND4: fast and accurate defocus estimation from electron micrographs. J Struct Biol 192: 216–221.

16. Lövestam S, Li D, Wagstaff JL, Kotecha A, Kimanius D, McLaughlin SH, Murzin AG, Freund SMV, Goedert M, Scheres SHW (2023) Disease-specific tau filaments assemble via polymorphic intermediates. BioRxiv.

17. Scheres SHW (2020) Amyloid structure determination in RELION-3.1. Acta Cryst D 76: 94–101.

18. Zivanov J, Otón J, Ke Z, von Kügelen A, Pyle E, Qu K, Morado D, Castaño-Diez D, Zanetti G, Bharat TAM et al (2022) A Bayesian approach to single-particle electron-tomography in RELION-4.0. eLife 11: e83724.

19. Scheres SHW, Chen S (2012) Prevention of overfitting in cryo-EM structure determination. Nature Meth 9: 853–854.

20. Emsley P, Lohkamp B, Scott WG, Cowtan K (2010) Features and development of Coot. Acta Crystallogr D 66: 486–501.

21. Yamashita K, Palmer CM, Burnley T, Murshudov GN (2021) Cryo-EM single particle structure refinement and map calculation using *Servalcat*. Acta Crystallogr D 77: 1282–1291.

22. Murshudov GN, Vagin AA, Dodson EJ (1997) Refinement of macromolecular structures by the maximum-likelihood method. Acta Crystallogr D 53: 240–255.

23. Murshudov GN, Skubák P, Lebedev AA, Pannu NS, Steiner RA, Nicholls RA, Winn MD, Long F, Vagin AA (2011) REFMAC fior the refinement of macromolecular crystal structures. Acta Crystallogr D 67: 355–367.

24. Chen VB, Arendall WB, Headd JJ, Keedy DA, Immormino RM, Ksaprai GJ, Murray LW, Richardson JS, Richardson DC (2010) MolProbity: all-atom structure validation for macromolecular crystallography. Acta Crystallogr D 66: 12–21.

25. Pettersen EF, Goddard TD, Huang CCX, Meng EC, Couch GS, Croll TI, Morris JH, Ferrin TE (2021) ChimeraX: structure visualization for researchers, educators, and developers. Protein Sci 30: 70–82.

26. Schrödinger L, DeLano W (2020) PyMol available at: <http//www.pymol.org/pymol>.

27. Stern AM, Yang Y, Jin S, Yamashita K, Meunier AL, Liu W, Cai Y, Ericsson M, Liu L, Goedert M et al (2023) Abundant Aβ fibrils in ultracentrifugal supernatants of aqueous extracts from Alzheimer’s disease brains. Neuron 111: 2012–2020.

28. Yang Y, Zhang W, Murzin AG, Schweighauser M, Huang M, Lövestam S, Peak-Chew SY, Saito T, Saido TC, Macdonald J et al (2023) Cryo-EM structures of amyloid-β filaments with the Arctic mutation (E22G) from human and mouse brains. Acta Neuropathol 145: 325–333.

29. Lövestam S, Schweighauser M, Matsubara T, Murayama S, Tomita T, Ando T, Hasegawa K, Yoshida M, Tarutani A, Hasegawa M et al (2021) Seeded assembly *in vitro* does not replicate the structures of α-synuclein filaments from multiple system atrophy. FEBS Open Bio 11: 999–1013.

30. Crooks EJ, Fu Z, Irizarry BA, Zhu X, Van Nostrand WE, Chowdhury S, Smith SO (2022) An electrostatic cluster guides Aβ fibril formation in cerebral amyloid angiopathy. BioRxiv.

31. Scheres SHW, Ryskeldi-Falcon B, Goedert M (2023) Molecular pathology of neurodegenerative diseases by cryo-EM of amyloids. Nature 621: 701–710.

32. Wagner J, Degenhardt K, Veit M, Louros N, Konstantoulea K, Skodras A, Wild K, Liu P, Obermüller U, Bansal V et al (2022) Medin co-aggregates with vascular amyloid-β in Alzheimer’s disease. Nature 612: 123–131.

33. Van Dyck CH, Swanson CJ, Aisen P, Bateman RJ, Chen C, Gee M, Kanekiyo M, Li D, Reyderman L, Cohen S et al (2023) Lecanemab in early Alzheimer’s disease. N Engl J Med 388: 9–21.

34. Sims JR, Zimmer JA, Evans CD, Lu M, Ardayfio P, Sparks JD, Wessels AM, Shcherbinin S, Wang H, Serap Monkul Nery E et al (2023) Donanemab in early symptomatic Alzheimer disease: the TRAILBLAZER-ALZ2 randomized clinical trial. JAMA 330: 512–527.

35. Jucker M, Walker LC (2023) Alzheimer’s disease: From immunotherapy to immunoprevention. Cell 186, 4260–4270.

